# Trophic microRNA: Precursor and mature microRNA ingestion downregulates the target transcripts and hampers larval development in *Plutella xylostella*

**DOI:** 10.1101/2023.12.21.572757

**Authors:** Rutwik Bardapurkar, Sagar Pandit

**Affiliations:** Agricultural Biotechnology and Chemical Ecology Research Laboratory, Department of Biology, Indian Institute of Science Education and Research, Pune, Maharashtra 411008, India

**Keywords:** Lepidoptera, let-7, *Plutella xylostella*, post-transcriptional gene regulation, trophic microRNA

## Abstract

MicroRNAs (miRNAs) are post-transcriptional gene regulators. In the miRNA pathway’s cytoplasmic part, the miRNA is processed from a hairpin-structured precursor (pre-miRNA) to a double-stranded (ds) mature RNA and ultimately to a single-stranded mature miRNA. In insects, ingesting these two ds forms can regulate the target gene expression; this inspired the trophic miRNA’s use as a functional genomics and pest management tool. However, systematic studies enabling comparisons of pre- and mature forms, dosages, administration times, and instar-wise effects on target transcripts and phenotypes, which can help develop a miRNA administration method, are unavailable due to the different focuses of the previous investigations. We investigated the impact of trophically delivered *Px*-let-7 miRNA on the Brassicaceae pest, *Plutella xylostella*, to compare the efficacies of its pre- and ds-mature forms. Continuous feeding on the miRNA-supplemented diet suppressed expressions of *FTZ-F1* and *E74*, the target ecdysone pathway genes. Both the pre-let-7 and mature let-7 miRNA forms similarly downregulated the target transcripts in all four larval instars. Pre-let-7 and let-7 ingestions decreased *P. xylostella*’s larval mass and instar duration and increased mortality in all instars, exhibiting adverse effects on larval growth and development. Pre-miRNA processing *Dicer-1* was upregulated upon pre-let-7 ingestion, indicating the miRNA uptake by the midgut cells. The scrambled sequence controls did not affect the target transcripts, suggesting the sequence-specific targeting by the mature miRNA and hairpin cassette’s non-involvement in the target downregulation. This work provides a framework for miRNA and target gene function analyses and potentiates the trophic miRNA’s utility in pest management.

## Introduction

The two major classes of small RNAs (smRNAs), microRNA (miRNA) and short-interfering RNA (siRNA), are involved in the post-transcriptional gene expression regulation (Kehl et al. 2017; Zhu and Palli 2020). miRNAs are transcribed from their genes, and siRNAs are generally the degradation products of long dsRNAs. Active forms of miRNAs, the 21-24 nt-long fragments (mature miRNAs or miRNAs), are formed by a three-step processing (Lucas and Raikhel 2013). Inside the nucleus, the long primary miRNA (pri-miRNA), a complex multiloop structure with 5’-cap and poly-A tail, is cleaved to produce a single stem-loop or hairpin structure called precursor miRNA (pre-miRNA) (Katahira and Yoneda 2011; Saito et al. 2005). The pre-miRNA then enters the cytoplasm, where Dicer-1 dices it to generate a 21-24 bp mature double-stranded (ds) miRNA; *Dicer-1*’s cleavage sites determine the generated miRNA duplex’s sequence (Zhu and Palli 2020). One of the strands is selected from the duplex and loaded onto Argonaute proteins (Czech et al. 2009). The RNA-induced silencing complex (RISC) transports this active miRNA strand to the complementary target mRNA to form the mRNA-miRNA duplex; the mRNA is then cleaved to effect the post-transcriptional gene regulation (Zhu and Palli 2020). Thus, in the miRNAs’ gene expression regulation site, the cytoplasm, only pre-, ds mature and mature forms are present.

The siRNAs originate from exogenously or endogenously produced long dsRNAs. *Dicer-2* dices the long dsRNAs into 18-24 nucleotide ds forms that cause the target transcript degradation with the help of RISC involving *AGO-2* (Vogel et al. 2019); this post-transcriptional gene silencing mechanism is known as RNA interference (RNAi). Since all the *Dicer-2*-generated fragments can bind to their complementary transcripts, there is a possibility of one long dsRNA causing RNAi in multiple genes. Such off-target effects can be troublesome when gene-specific silencing is desired. In contrast, pre-miRNAs produce only one or few target-specific mature miRNAs; thus, the miRNA-based gene regulation has evolved to be target transcript-specific (Kehl et al. 2017)

Usually, the smRNAs cause post-transcriptional gene regulations within their producer individual’s cells. Several studies revealed that many smRNAs, especially miRNAs, are induced in plants upon herbivory (Bozorov et al. 2012; Jeyaraj et al. 2017). Recent reports also showed that the ingested dsRNA, especially the plant-origin dsRNA, causes post-transcriptional gene silencing in herbivore insects (Kumar et al. 2012; Kumar et al. 2014; Mao et al. 2007). When collectively considered, these reports provided an origin for the hypothesis that hostplant’s smRNAs regulate insect herbivores’ target gene expression upon their co-ingestion with the plant (Zhang et al. 2019). In insects, especially in lepidopterans, where the occurrence of crucial siRNA pathway components like the RNA-dependent RNA polymerase (RdRP) is still unconfirmed (Mamta and Rajam 2017; Tomoyasu et al. 2008), the miRNA pathway machinery is complete and well-characterized (Doench et al. 2003; Gordon and Waterhouse 2007). In these insects, the miRNA pathway apparatus processes the exogenously acquired siRNAs (Doench et al. 2003). miRNA genes, targets, and miRNA pathway components are conserved, suggesting that the miRNA pathway could be the evolutionarily selected smRNA pathway in these insects (Obbard et al. 2006). Moreover, the precursor hairpin structures are more stable than the other dsRNA molecules (Bonnet et al. 2004). Therefore, there is a growing interest in testing miRNAs as pesticide candidates. Moreover, since the miRNAs have evolved to be target-specific, the cross-kingdom activity is surmised to be a naturally evolved miRNA-based mechanism in plant-insect interactions (Zhang et al. 2019). Efforts to test these hypotheses showed that, like siRNAs, ingested miRNAs also regulated insect target genes (Etebari et al. 2018). Moreover, they showed phenotypic effects in many insects. In *Spodoptera exigua* Hübner (Lepidoptera: Noctuidae), they hampered larval growth and affected molting (Zhang et al. 2015). In *Helicoverpa armigera* Hübner (Lepidoptera: Noctuidae), they increased larval and adult mortality (Saini et al. 2018). In *Bombyx mori* L. (Lepidoptera: Bombycidae), several miRNAs with specific temporal and spatial expressions regulated larval development, pupal maturation, and adult emergence (Yu et al. 2008). Ingested miR-34-5p targeting the ecdysone receptor caused high mortality, low fecundity, and developmental defects in three insect species (Li et al. 2023). Similar effects were demonstrated using pre-miRNAs with engineered target binding cassettes overexpressed in plants targeting the *H. armigera*’s acetylcholinesterase transcripts (Bally et al. 2020).

Indeed, the findings of the above studies were promising and indicated the active roles of ingested miRNAs (Kolliopoulou and Swevers 2014; Kumar et al. 2022). Since insects ingest high levels of cytoplasmic forms of plant miRNAs [pre-ds mature, and mature] during herbivory, it is crucial to understand whether these forms can function in the insect when trophically acquired. The dsRNAs are highly stable in the gut lumen compared to their highly labile single-stranded forms (Zhang et al. 2021). Therefore, mature miRNAs’ ds forms are considered ‘active forms’ in the trophically acquired RNA pool; consequently, these ds forms are often used in trophic delivery studies (Li et al. 2023; Zafar et al. 2021). Functioning of such trophically delivered pre- and ds mature miRNAs will require 1) their uptake in the gut cells from the gut lumen, 2) processing of pre- and ds-forms to the mature ones, 3) strand selection, and 4) transport and binding to the target transcript. A comprehensive study covering all these aspects is desired. We conducted a study comparing the effects of pre- and mature ds conformations. We used Brassicaceae crops’ serious pest *Plutella xylostella* (L.) (Lepidoptera: Plutellidae) as a model (Fathipour and Mirhosseini 2017; Li et al. 2021; Zalucki et al. 2012). Besides being an agro-economic threat, *P. xylostella* showed the presence of plant-derived miRNAs in the larval hemolymph (Zhang et al. 2019), indicating an uptake of dietary miRNAs and posing it as a promising model system to understand the effects of ingested pre- and mature-miRNAs. We used *Px*-let-7 as a model miRNA (Liang et al. 2013). let-7 is conserved in insects (Lee et al. 2016; Roush and Slack 2008). In *B. mori*, it regulates the ecdysone pathway regulators *FTZ-F1* and *E74*, and its downregulation shows larval-larval and larval-pupal development arrests (Ling et al. 2014; Song and Zhou 2020). As it is highly conserved and plays a vital role in insect development, we used *P. xylostella*’s let-7 to systematically characterize the effect of its pre- and mature ds forms’ ingestions on 1) the endogenous pre- and mature let-7 levels, 2) the target *Px-FTZ-F1* and *Px-E74* transcripts, 3) larval development, and 4) larval mortality in all four instars (Fig. 1). *P*. *xylostella*’s smRNA (miRNA and RNAi) pathways are understudied. Especially their responses to the exogenous smRNAs are not known. Therefore, we also analyzed whether these pathways’ components’ transcripts indicate the pre- and mature-miRNA intake.

**Fig. 1.**
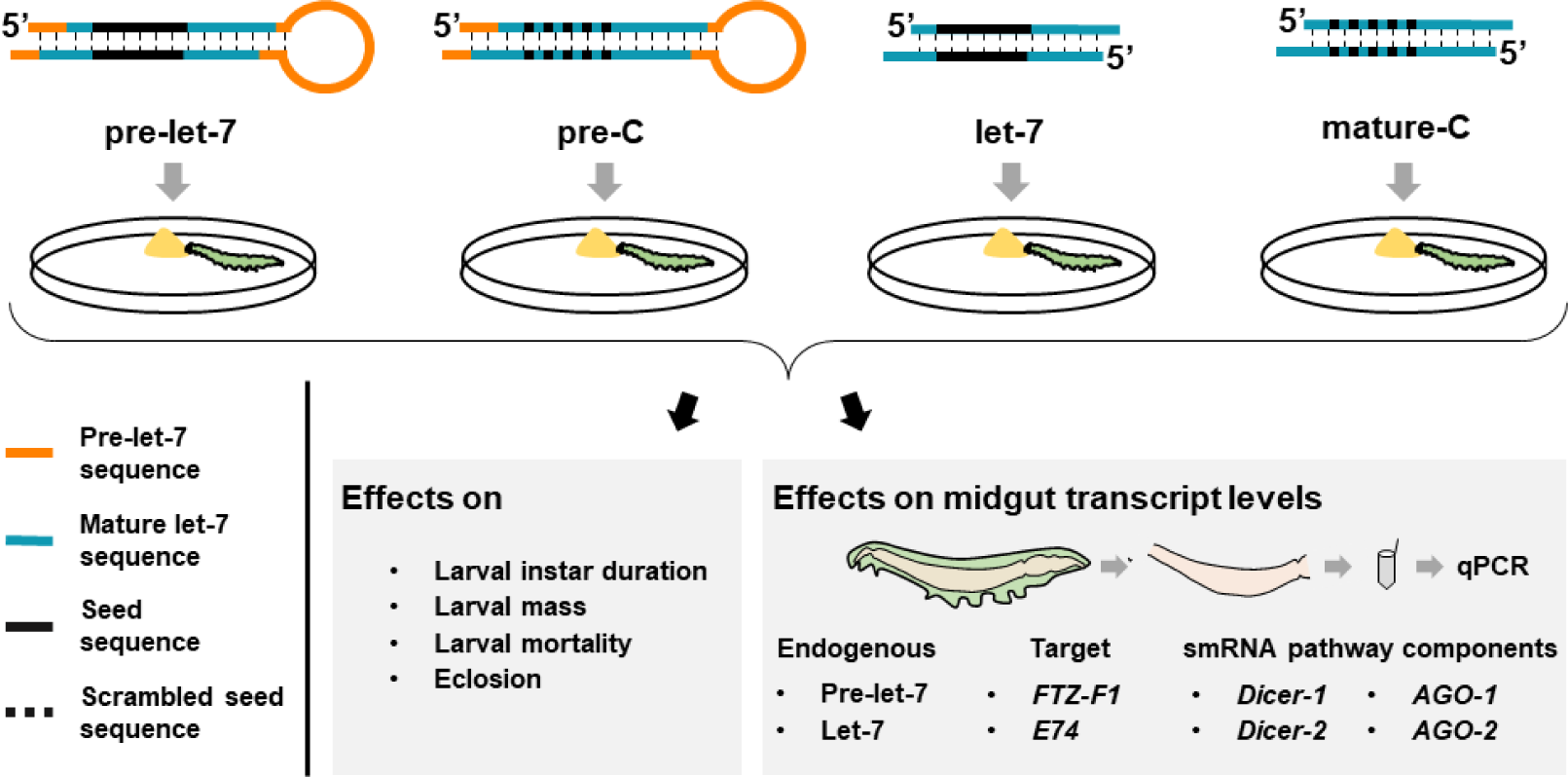
A schematic of the experimental design, miRNA and respective control’s structural details, feeding assay setup, analyzed transcripts and phenotypes.

## Materials and methods

### Insects

Three generations of *Plutella xylostella* were raised by feeding the larvae on an artificial diet (AD) at 24 °C, 70 % relative humidity, and a 16:8 h light: dark photoperiod, as described by Shelton et al. (1991). Insects of the fourth and later generations were used in all the experiments. The same AD was used in all the experiments.

### RNA isolation, cDNA synthesis, and qPCR

Five biological replicates, each containing midguts of 25 larvae fed on their respective diets, were used to analyze transcript levels. Larvae were dissected, and their midguts were thoroughly washed in the ice-cold Tris-EDTA buffer to remove the sticking diet, pooled into 250 µl RNAiso Plus reagent (Takara, Japan), and kept at 4 °C until use. Total RNA was isolated following the manufacturer’s instructions and precipitated overnight at −20 °C using pre-chilled 2.5 volumes of isopropanol to ensure the smRNA recovery. All mRNA and pre-let-7 samples’ cDNAs were synthesized using 100 ng RNA and the PrimeScript Reverse Transcriptase kit (Takara, Japan). As previously described (Chen et al. 2005), the let-7 and 5S rRNA cDNAs were independently synthesized from the total RNA using stem-loop primers (5 nM). qPCRs were performed using the SYBR Premix Ex Taq II reagent kit (Takara, Japan) on the CFX96 Touch Real-Time PCR Detection System (Biorad, USA).

Thermocycling consisted an initial denaturation step of 95 °C for one min, followed by 44 cycles of 95 °C for 45 s, 60 °C for 45 s, and 72 °C for 45 s; Ubiquitin was used as an internal reference for the relative mRNA and pre-let-7 quantitation, and 5S rRNA for let-7 quantitation. Oligos used for cDNA synthesis, and qPCRs are given in Table S1.

### Let-7 expression dynamics during the larval development

To understand the let-7 expression dynamics during larval development, we analyzed the pre-let-7 and let-7 levels in the early (5 h after hatching or previous molting), mid (30 h after hatching or previous molting), and late (60 h after hatching or previous molting) stages of the larval instars.

### miRNA synthesis

Pre-let-7 and let-7 were synthesized using the DNA templates with upstream T7 promoter sequences (Table S2). Four forms: pre-let-7, its negative control (pre-C; a pre-let-7 cassette with a scrambled seed sequence in the hairpin such that the arising mature form has no predicted target in the *P. xylostella* gene pool), mature ds let-7 (hereafter termed let-7), and its negative control (mature-C; mature ds let-7 with the same scrambled seed sequence, as that in pre-C) (Fig. 1; Table S3). To ensure its stability in the larval gut lumen, we used the ds forms of let-7 and mature-C in the feeding experiments. The two dsRNA strands were independently transcribed, reassociated to form the mature duplexes, and stored at −20 °C until further use. *In vitro* transcription reactions were performed, as previously described by Donze and Picard (2002). In each 50 µl reaction, a 200 pmol DNA template was incubated with 1 mM rNTPs, 40 mM Tris–HCl pH 7.9, 6 mM MgCl2, 10 mM DTT, 10 mM NaCl, and 2 mM spermidine, 100 U T7 polymerase, and 40 U recombinant Ribonuclease inhibitor (Takara, Japan). Reactions were incubated at 37 °C for 4 h and then treated with 1 U DNAse (Himedia, India). Sense and antisense strands were reassociated by slow cooling and purified, as per Donze and Picard (2002). These miRNAs were hand-mixed with AD for the trophic delivery experiments. Since the exogenous miRNAs absorbed from the environment (in this case, gut lumen) act in the cytoplasm (the gene expression regulation cite), we used only the cytoplasm-specific ds forms (pre- and mature-) for feeding the larvae.

### miRNA concentration optimization for the trophic delivery

Using the first instar larvae, we determined the pre- and mature let-7’s dietary concentrations affecting the target transcripts, instar duration (hatching to molting), larval mass (40 h after hatching), and mortality (40 h after hatching). 4, 7, 10, 13, and 16 µg (in 50 µl DEPC-treated water) pre-let-7 or let-7 were hand-mixed per gram AD A separate assay was conducted for every treatment and control in 20ml polypropylene containers. Five larvae were kept in each assay container with *ad libitum* feeding; larvae fed on AD mixed with 50 µl DEPC-treated water were used as controls (C). Larvae were transferred to the fresh vials containing fresh diets every 24 h to avoid miRNA degradation.

### miRNA stability in AD

Using qPCR, we verified whether the pre-let-7 and let-7 miRNAs were stable in the AD during the feeding period. Since we provided the larvae with fresh diets every 24 h in all the experiments, we analyzed the miRNA stability in AD for 24 h. Total RNA was isolated from the AD containing 0 and 10 µg/g RNA after 0 and 24 h of mixing. For collecting the 24 h samples, fourth instar larvae were fed on these ADs for 24 h (as described above) to ensure that they received the same larval activity and excreta as the assay diets. RNA isolation (from 100 µg samples), cDNA synthesis, and qPCR were conducted as detailed above. To quantify pre-let-7, 100 ng synthetic ubiquitin RNA (Table S4) was mixed with each sample collected in 250 µl RNAiso Plus reagent. Similarly, 100 ng synthetic ds 5s rRNA (Table S4) was mixed with each let-7 quantification sample. Five replicates (n= 5) per every treatment and control were used.

### miRNA feeding duration optimization

We analyzed the temporal changes in the target transcripts *FTZ-F1* and *E74* to determine the miRNA feeding duration required for their effective downregulation. We compared the midgut *FTZ-F1* and *E74* transcript levels of the control and miRNA-complemented diet-fed first instar larvae of 5, 15, 30, 45, and 60 h ages. At 24°C, the molting process begins around 70 h in all four instars (pupation in the fourth instar); to ensure the sampling of actively feeding larvae and avoid the molting process-entered physiologically differed contaminants, the larvae of >60 h age were not used in any of the experiments.

We also analyzed the transcripts of two miRNA pathway genes, *Dicer-1* and *AGO-1*, and two siRNA pathway genes, *Dicer-2* and *AGO-2*, in the miRNA-complemented diet-fed first instar larvae of 5, 15, 30, 45, and 60 h ages, to assess whether target gene-regulation and smRNA pathway components’ expression dynamics’ timings coincide.

### Effects of miRNA ingestion on the endogenous pre-let-7 and let-7 levels

To verify that the pre-let-7 and let-7 ingestion increased their levels in the midgut cells, we assessed the larvae’s pre- and mature let-7 levels fed on all five treatments. Every instar’s 60-h-old larvae were used for this analysis. Levels in the miRNA-ingested larvae were compared to the ones in the control larvae.

### Effects of miRNA ingestion on target transcripts

To test whether the ingested pre- and mature let-7 affect the target transcript levels in all four larval instars, we analyzed let-7’s target transcripts *FTZ-F1* and *E74* in the larvae fed on different miRNA-containing and control diets. Every instar’s 60 h-old larvae were used for this analysis.

### Effects of pre- and mature let-7 feeding on *P. xylostella* growth and development

To determine the impacts of pre- and mature let-7 ingestions on *P. xylostella* performance, we analyzed the larval instar durations, mass, mortality (%), and eclosion (%). Larvae were fed the pre-let-7 and let-7-supplemented diets from the neonate stage. Mass and mortality were measured in the 60-h-old larvae of every instar. In the instar duration, larval mass, mortality, and eclosion analyses, 30 larvae (n= 30) were used per every treatment and control.

### Dynamics of the small RNA pathway components’ transcripts upon let-7 ingestion

We analyzed the transcripts of the smRNA pathway components in the midguts of every instar’s 60-h-old larvae fed on various treatment and control ADs to understand their instar-wise responses to the ingested miRNAs.

### Data analysis

Fold changes between two means, *a* and *b,* were calculated as 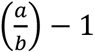, where a> b. Levene’s test was used to assess the homogeneity, and the Jarque-Bera test was used to assess the normality of the quantitative data (mean± SE). Parametric single-factor data were analyzed using one-way ANOVA; Tukey’s *post hoc* test was used to determine the statistical significance (*p*≤ 0.05). Non-parametric single-factor data were analyzed using the Kruskal-Wallis test and Dunn’s *post hoc* test (*p*≤ 0.05). Non-parametric two-or multi-factor data were analyzed by Friedman’s two-way ANOVA and Dunn’s *post hoc* test (*p*≤ 0.05). All the statistical analyses were conducted using the IBM SPSS Statistics. Directions and strengths of correlations between miRNA levels and transcript levels of miRNA pathway and target genes were calculated (separately for each larval instar) using the non-parametric Spearman’s Rho (*r_s_*) tests, and the significances (*p*≤ 0.05) were determined using a 2-tailed test.

## Results

### let-7 expression dynamics during the larval development

Pre-let-7 levels varied during all four larval instars’ early, mid, and late (pre-molting) stages (Fig. S1). In the late (pre-molting) stages of the first, second, third, and fourth instars, the pre-let-7 levels increased by ≥7.6-fold (*p*≤ 0.03) (χ^2^= 10.5, *p=* 0.005), ≥6.6-fold (*p=* 0.03) (χ^2^= 10.5, *p=* 0.005), ≥4.3-fold (*p=* 0.04) (χ^2^= 9.62, *p=* 0.008), and ≥7.3-fold (*p=* 0.01) (χ^2^= 9.62, *p=* 0.008), respectively, than the respective early and mid-stages (Fig. S1a). Let-7 also showed the same profile (Fig S1b). The early and mid-stage pre-let-7 and let-7 levels differed in none of the instars.

### miRNA concentration optimization for the trophic delivery

Pre-let-7 and let-7 mixed with AD did not degrade in the 24 hours, after which the larvae were provided freshly complemented diets (Fig. S2). Transcript levels of both the targets *FTZ-F1* (Fig. 2a) and *E74* (Fig. 2b) dropped at the 10 µg/g AD concentrations of pre-let-7 and let-7; they remained unchanged at pre-let-7 and let-7 concentrations higher than 10 µg/g. The *FTZ-F1* transcripts showed an 18-fold (*p=* 0.02) (χ^2^_(11)_= 41.06, *p<* 0.001) decrease upon the ingestion of 10 µg/g and higher pre-let-7 concentrations, compared to the C larvae (χ^2^_(11)_= 44.72, *p<* 0.0001). With let-7, *FTZ-F1* transcripts showed more severe suppression at 10 µg/g and higher concentrations; they showed a >40-fold (*p=* 0.002) decrease than the controls. Likewise, the *E74* transcript levels showed >22-fold (*p=* 0.04) reduction on the ≥10µg/g concentration pre-let-7 diets and >33-fold (*p=* 0.006) reduction on the ≥10µg/g concentrations let-7 diets than C (χ^2^_(11)_= 43.68, *p<* 0.0001).

**Fig. 2.**
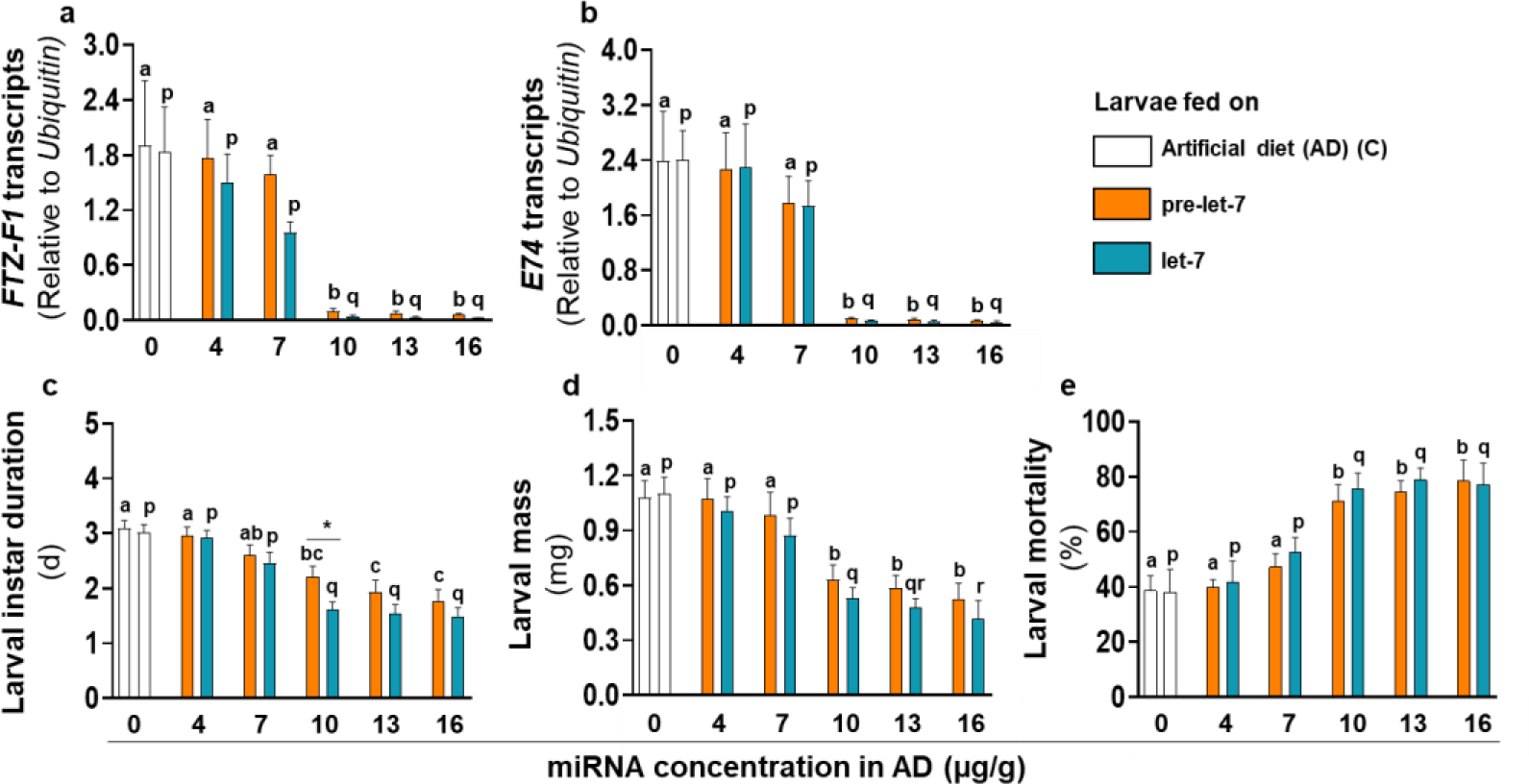
miRNA concentration optimization for the trophic delivery in the first instar *P. xylostella* larvae. **a** *FTZ-F1* and **b** *E74* transcripts **c** Larval instar duration (mean± SE, n= 30 larvae), **d** larval mass (mg) (mean± SE, n= 30), and **e** larval mortality after 24 h (%) (n= 5 sets, each containing 30 larvae). Significant differences were determined using Friedman’s two-way ANOVA and Dunn’s *post hoc* test (*p≤* 0.05). Letters a, b, and c indicate significant differences between pre-let-7-fed larvae, and letters p, q, and r indicate significant differences between let-7-fed larvae. Asterisk indicates the significant difference between pre-let-7 and let-7-fed larvae.

Similar effects were observed on the first instar duration (Fig. 2c), larval mass (Fig. 2d), and mortality (Fig. 2e). The first instar duration decreased by >0.3-fold (*p=* 0.004), >0.5-fold (*p<* 0.0001*=*), and >0.5-fold (*p=* 0.000) at the 10, 13, and 16 µg/g pre-let-7 concentrations, respectively than the control (χ^2^_(11)_= 103.94, *p<* 0.001). On the let-7-complemented diet, the first instar duration showed a ≥ 0.8-fold decrease at 10, 13, and 16µg/g concentrations (*p<* 0.0001) than the control. The first instar duration was ≥1.1-fold (*p=* 0.02) lower on the 10 µg/g let-7 diet than on the 10 µg/g pre-let-7 diet.

Both pre-let-7 and let-7 diets showed similar effects on larval mass across their tested concentrations (Fig. 2d). Larval mass was ≥0.7-fold lower on the 10, 13, and 16 µg/g (*p<* 0.001) pre-let-7 diets and ≥1.0-fold lower (*p<* 0.001) on the 10, 13 and 16µg/g let-7, diets than the respective controls (χ^2^_(11)_= 130.2, *p<* 0.001).

pre-let-7 and let-7 ingestion also affected the larval mortality (Fig. 2e). At 10, 13, and 16 µg/g concentrations, the larval mortality increased by ≥0.8-fold (*p=* 0.01) on the pre-let-7 diet and by ≥0.9-fold (*p=* 0.01) on the let-7 diet (χ^2^_(11)_ = 37.49, *p<* 0.001) than the respective controls. For both the miRNA conformations, larval mortality did not change with the higher miRNA concentrations than 10 µg/g.

Ingestion of both pre- and mature forms showed significant effects on the target transcripts, instar duration (hatching to the first molting), larval mass, and mortality upon 10 µg/g concentration ingestion; these effects saturated at the higher miRNA concentrations than 10 µg/g. Therefore, we used 10 µg/g miRNA concentrations of pre-let-7 and let-7 in the further experiments.

### miRNA feeding duration optimization

The endogenous target transcript levels did not vary with the larval age [*FTZ-F1*: χ^2^_(14)_= 39.4, *p=* 0.003 (Fig 3a); *E74*: χ^2^_(14)_= 44.3, *p<* 0.0001 (Fig. 3b)]. Upon continuous pre-let-7 and let-7 ingestions for 45 h, target transcripts reduced, compared to the C larvae. *FTZ-F1* showed 0.9-fold (*p=* 0.04) and 2.5-fold (*p=* 0.009) reductions than C after 45h of pre-let-7 and let-7 ingestions, respectively. After 60 h, these levels further decreased by 12.3-fold (*p<* 0.001) and 15.6-fold (*p<* 0.0001) than C. Similarly, *E74* levels dropped by 0.9-fold (*p=* 0.01) and 3.1-fold (*p=* 0.001) after 45 h feeding on pre-let-7 and let-7 diets, respectively, compared to C. After 60 h of pre-let-7 and let-7 ingestion, these levels decreased by 13.7-fold (*p<* 0.001) and 35.2-fold (*p<* 0.001) than the endogenous (C) levels, respectively. When the different miRNA ingestion durations were compared, levels of the target *FTZ-F1* and *E74* transcripts after 60 h ingestion of both pre-let-7 and let-7 were ≥7.6-fold lower (*p<* 0.02) than the larvae ingesting these respective miRNAs for 5, 15, and 30 h; 45 h transcript levels did not differ from the respective shorter (5, 15, and 30 h) and longer (60 h) miRNA ingestion duration levels.

**Fig. 3.**
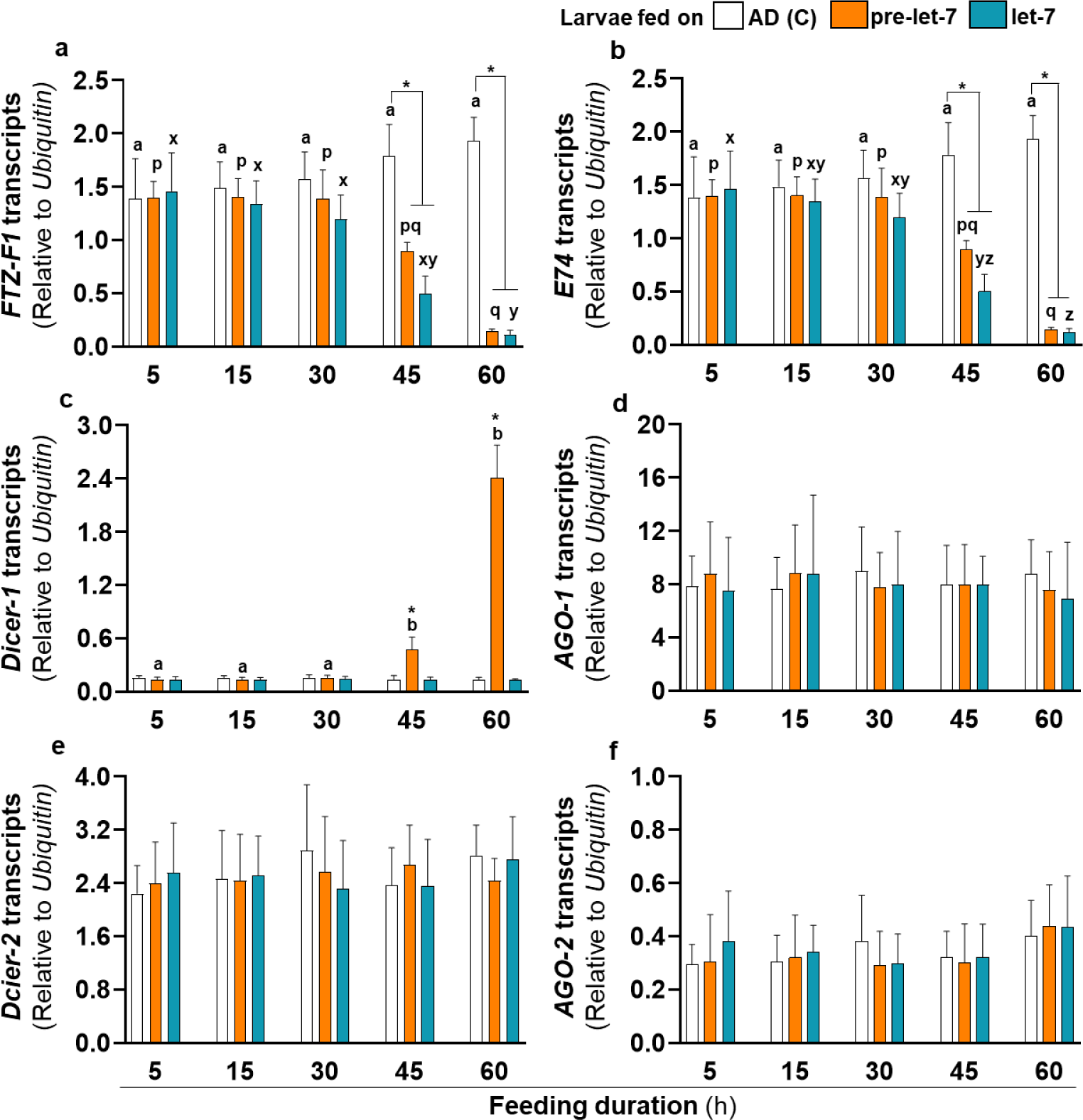
miRNA feeding duration optimization. Transcripts of target **a** *FTZ-F1* and **b** *E74*, miRNA pathway genes **c** *Dicer-1*, **d** AGO-1, and siRNA pathway genes **e** *Dicer-2* and **f** AGO-2. Levels (relative to *Ubiquitin*) are from larval midgut (n= 5 RNA samples, each containing 25 larvae’s midguts). Significant differences were determined using Friedman’s two-way ANOVA and Dunn’s *post hoc* test (*p≤* 0.05). Letters a, b, and c indicate significant differences between the controls (endogenous transcript levels); letters p, q, and r indicate significant differences between the pre-let-7-fed larvae of different instars; and letters x, y, and z indicate significant differences between the let-7-fed larvae of different instars. Asterisks indicate significant differences between the treatment groups of one feeding duration.

Like the target transcripts, *Dicer-1* transcript levels also varied only after 45 h (Fig. 3c). Among all the treatments and controls, they varied only after the pre-let-7 ingestion. After 45 h of pre-let-7 feeding, the *Dicer-1* levels increased by 2.6-fold (*p*= 0.01), 2.5-fold (*p*= 0.02), and 2.1-fold (*p*= 0.04), compared to 5, 15, and 30 h, respectively; they further increased by 17.2-fold (*p*= 0.003), 16.8-fold (*p*= 0.004), and 14.8-fold (*p*= 0.010) after 60 h, compared to 5, 15, and 30 h, respectively (χ^2^_(14)_= 25.33, *p*= 0.03). At both 45 and 60 h, *Dicer-1* levels were 2.5-fold (*p*= 0.005) and 17.5-fold (*p*=0.003) higher, respectively, than the C larvae; they were 2.6-fold (*p*= 0.01) and 17.7-fold (*p*=0.006) higher than the let-7 feeding larvae at 45 and 60 h, respectively. Transcripts of the other miRNA pathway gene, *AGO-1* (Fig. 3d), and siRNA pathway genes *Dicer-2* (Fig. 3e) and *AGO-2* (Fig. 3f) did not vary in any of the treatments and controls.

### Effects of miRNA ingestion on the endogenous pre-let-7 and let-7 levels

Pre-let-7 ingested larvae showed increase in the midgut pre-let-7 levels compared to the C, pre-C and mature-C controls of all four instars (Fig. 4a). The first instar showed a 1.5-fold increase (*p*< 0.04) (χ^2^= 4.35, *p=* 0.01), and the second instar showed a ≥2.0-fold increase (*p*≤ 0.01) (χ^2^= 10.64, *p*≤ 0.008) in pre-let-7 levels, compared to their respective controls. The third instar did not show a significant pre-let-7 induction (χ^2^= 7.23, p =0.1). The fourth instar showed a 2.3-fold increase (*p=* 0.04) (*F*_4,20_= 3.54, *p=* 0.02) in pre-let-7 levels compared to mature-C; although it also showed a 2.1-fold increase compared to C and pre-C, it was marginally non-significant (*p=* 0.05).

**Fig. 4.**
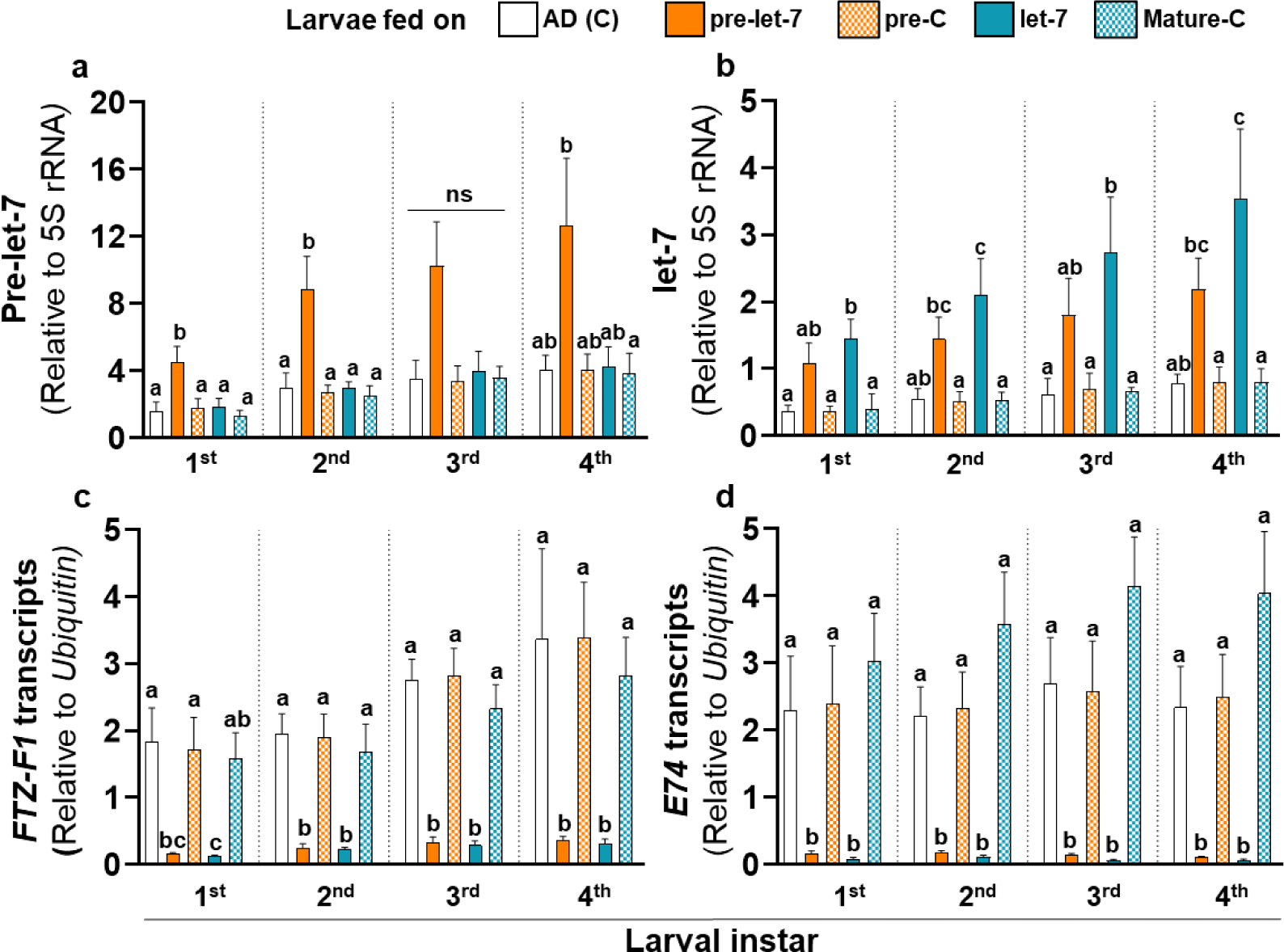
Effects of pre-let-7 and let-7 ingestion on various transcripts in different larval instars. Levels of **a** pre-let-7 (relative to *Ubiquitin*), **b** let-7 (relative to 5s rRNA), **c** *FTZ-F1* (relative to *Ubiquitin*), and **d** *E74* (relative to *Ubiquitin*) in the midguts of larvae feeding on C, pre-let-7, pre-C, let-7, and mature-C diets (n= 5 RNA samples, each containing 25 larvae’s midguts). Significance in **a** (first instar and fourth instar) and **b** (first instar) was determined by one-way ANOVA and Tukey’s *post hoc* test (*p≤* 0.05). Significance in all other comparisons was determined by the Kruskal-Wallis test and Dunn’s *post hoc* test (*p≤* 0.05). Letters a, b, and c denote differences between the larvae of the same instar feeding on different diets.

Let-7 levels increased only on the let-7 diets compared to the C, pre-C and mature-C ones (Fig. 4b). They were 2.6-fold higher (*p*≤ 0.02) (χ^2^= 4.82, *p=* 0.006) in the first instar, >2.9-fold higher (*p*≤ 0.02) (χ^2^= 11.62, *p<* 0.02) in the second instar, >2.9-fold higher (*p*≤ 0.02) (χ^2^= 10.18, *p* = 0.03) in the third instar, and ≥3.4-fold higher (*p*≤ 0.018) (χ^2^= 12.92, *p=* 0.01) in the fourth instar than C, pre-C and mature-C controls.

### Effects of miRNA ingestion on target transcripts

We tested the effects of pre-let-7 and let-7 ingestion on the target genes’ transcripts in all four instars. *FTZ-F1* transcript levels were 10.8-fold (*p=* 0.007) (χ^2=^ 17.41, *p=* 0.001), 6.7-fold (*p=* 0.004) (χ^2=^ 17.4, *p=* 0.001), 7.5-fold (*p=* 0.006) (χ^2=^ 17.8, *p=* 0.001) and 8.4-fold (*p=* 0.01) (χ^2=^ 17.42, *p=* 0.001) lower in the pre-let-7-ingested first, second, third, and fourth instars, respectively, than their respective controls (Fig. 4c). Similarly, *FTZ-F1* transcript levels were 14.1-fold (*p=* 0.003), 7.3-fold (*p=* 0.005), 8.7-fold (*p=* 0.004), and 9.6-fold (*p=* 0.007) lower in the let-7-ingested first, second, third, and fourth instars, respectively, than their respective controls. *FTZ-F1* transcripts showed strong negative correlations with the let-7 levels in all four instars (first instar: *r*= −0.96, *p=* 0.007; second instar: *r*= −0.94, *p=* 0.01; third instar: *r*= −0.92, *p=* 0.02; fourth instar: *r*= −0.91, *p=* 0.02).

*E74* transcript levels were 12.9-fold (*p=* 0.03) (χ^2=^ 18.26, *p=* 0.001), 11.6-fold (*p=* 0.03) (χ^2=^ 18.57, *p*< 0.0001), 18.4-fold (*p=* 0.04) (χ^2=^ 18.78, *p*< 0.0001) and 20.2-fold (*p=* 0.03) (χ^2=^ 19.59, *p*< 0.0001) lower in the pre-let-7-ingested first, second, third, and fourth instars, respectively, than their respective controls (Fig. 4d). These levels were 30.4-fold (*p=* 0.003), 18.6-fold (*p=* 0.008), 45.0-fold (*p=* 0.005), and 34.9-fold (*p=* 0.003) lower in the let-7-ingested first, second, third, and fourth instars, respectively, than their respective controls. The target genes’ transcripts showed no level change in the larvae fed on pre-let-7-control (pre-C), let-7-control (mature-C), and C diets. *E74* transcripts showed strong negative correlations with the let-7 levels in the first three instars (first instar: *r*= −0.94, *p=* 0.01; second instar: *r*= −0.88, *p=* 0.04; third instar: *r*= −0.88, *p=* 0.04). The fourth instar also showed a strong negative correlation (*r*= −0.85); however, it was marginally non-significant (*p=* 0.06).

### Ingested pre-let-7 and let-7’s effects on *P. xylostella* growth and development

Pre-let-7 and let-7-ingestion reduced the durations of all four instars as compared to the three controls C, pre-C, and mature-C (Fig. 5a). The instar-wise duration reductions in the pre-let-7 and let-7-fed larvae were as follows-first instar: ≥0.3-fold (*p*≤ 0.001) and ≥0.8-fold (*p<* 0.0001), respectively (*F*_4,145_= 19.45, *p<* 0.0001); second instar: ≥0.2-fold (*p<* 0.02) and ≥0.5-fold (*p<* 0.0001), respectively (*F*_4,145_= 11.56, *p<* 0.0001); third instar: ≥0.2-fold (*p<* 0.07) and ≥0.3-fold (*p<* 0.0001), respectively (*F*_4,145_= 6.35, *p<* 0.0001); and fourth instar:≥0.3-fold (*p<* 0.002) and ≥0.5-fold (*p<* 0.0001), respectively (χ^2=^ 37.85, *p<* 0.0001).

**Fig. 5.**
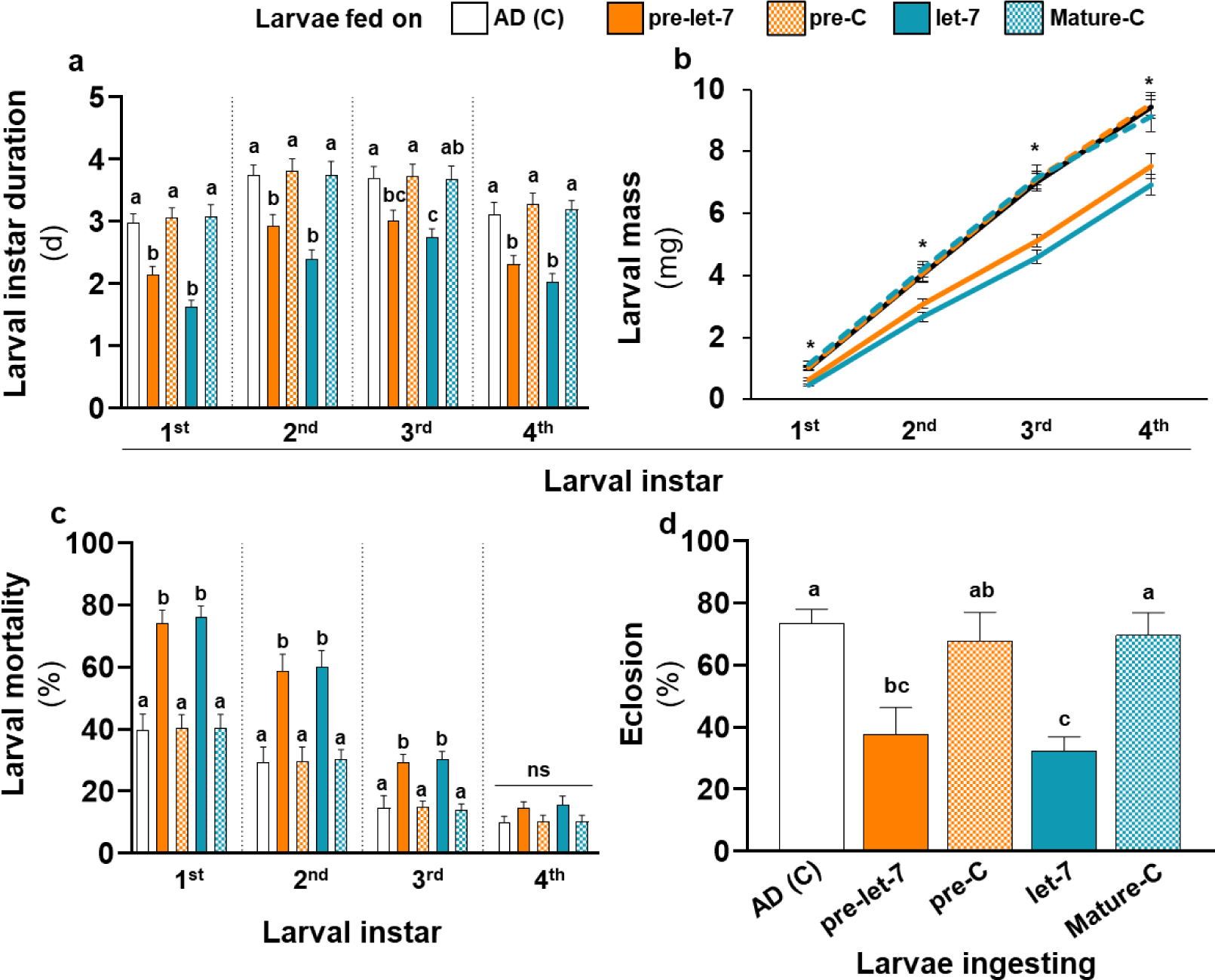
Effects of pre-let-7 and let-7 ingestion on growth and development. **a** Larval instar duration (mean± SE, n= 30), **b** larval mass (mg) (mean± SE, n= 30), **c** larval mortality (%) (n= 5 sets, each containing 30 larvae), and **d** pupal eclosion (%) (n= 5 sets, each containing 100 pupae) for each instar. In **a (**first to third instar), (**c**), and (**d**), significant differences were determined using one-way ANOVA and Tukey’s *post hoc* test (*p*≤ 0.05) and in **a** (fourth instar), and (**b**), using the Kruskal-Wallis test and Dunn’s *post hoc* test (*p*≤ 0.05). Letters a, b, and c in (**a**), (**c**), and (**d**), and asterisks in (**b**) denote differences between the larvae of the same instar feeding on different diets.

Pre-let-7- and let-7 ingestion also showed negative effects on the larval mass in all four instar stages as compared to the three controls (Fig. 5b). The instar-wise mass reductions in the pre-let-7 and let-7-fed larvae were as follows-first instar: ≥0.5-fold (*p*≤ 0.0002) and ≥1.1-fold (*p<* 0.0001), respectively (χ^2=^ 54.39, *p<* 0.0001); second instar: ≥0.3-fold (*p*≤ 0.0004) and ≥0.5-fold (*p<* 0.0001), respectively (χ^2=^ 31.83, *p<* 0.0001); third instar: ≥0.3-fold (*p<* 0.0001) and ≥0.5-fold (*p<* 0.0001), respectively (χ^2=^ 58.39, *p<* 0.0001); and fourth instar: ≥0.3-fold (*p*≤ 0.0001) and ≥0.3-fold (*p<* 0.0001), respectively (χ^2=^ 31.13, *p<* 0.0001). The masses of pre-let-7 and let-7 feeding larvae did not differ in any instars. The masses of pre-C or mature-C ingesting larvae did not differ from the no-RNA controls.

Mortality increased in the larvae ingesting pre-let-7 and let-7 (Fig. 5c). Compared to the controls, the mortality of pre-let-7 and let-7 ingesting first instar larvae increased by ≥0.8-fold (*p*< 0.0001) and ≥0.9-fold (*p<* 0.0001), respectively (*F*_4,20_= 18.45, *p<* 0.0001), in the second instar, by ≥1.0-fold (*p*< 0.0001) and ≥1.0-fold (*p<* 0.0001), respectively (*F*_4,20_= 11.4, *p<* 0.0001), and in the third instar, by ≥1.0-fold (*p*≤ 0.001) and ≥1.0-fold (*p<* 0.0001), respectively (*F*_4,20_= 8.88, *p<* 0.0001). On both pre-let-7 and let-7 diets, mortality of the fourth instars did not differ from the controls. All the survived fourth instar larvae pupated; pupation (%) was not different in treatments and controls.

pre-let-7 and let-7 ingesting larvae’s pupae showed ≥0.9-fold (*p*≤ 0.01) and ≥1.2-fold (*p*≤ 0.004) lower eclosion, respectively, than the controls (*F*_4,20_= 7.42, *p<* 0.0001) (Fig. 5d).

### Dynamics of the small RNA pathway components’ transcripts upon let-7 ingestion

*Dicer-1* showed upregulation upon pre-let-7 ingestion (Fig. 6a). The first and fourth instars showed a ≥16.6-fold (*p*≤ 0.005) (χ^2=^ 11.79, *p=* 0.019) and ≥9.2-fold (*p*≤ 0.028) (χ^2=^ 13.85, *p=* 0.007) upregulation, respectively than their respective controls. In the second instar of the pre-let-7-fed larvae, the *Dicer-1* levels were12.6-fold higher than pre-C (*p=* 0.01), 24.3-fold higher than mature-C (*p*< 0.0001), and 16.6-fold higher than let-7-fed larvae (*p=* 0.004) (χ^2=^ 14.61, *p=* 0.005); pre-let-7-fed second instar larvae’s *Dicer-1* levels were not different from the C larvae (p*=* 0.07); however, they were >12.0-fold (p≤ 0.01) higher than the other two controls. The pre-let-7 feeding third instars showed 33.0-fold (*p=* 0.001) upregulation than mature-C and 22.7-fold (p< 0.0001) upregulation than let-7-fed larvae (χ2= 18.09, p*=* 0.001). pre-let-7-fed third instar larvae’s *Dicer-1* levels were not different from the pre-C and C larvae. *Dicer-1* transcripts showed strong positive correlations with the pre-let-7 levels in all four instars (first instar: *r*= 0.98, *p=* 0.002; second instar: *r*= 0.99, *p*< 0.0001; third instar: *r*= 0.98, *p=* 0.001; fourth instar: *r*= 0.99, *p=* 0.0002).

**Fig. 6.**
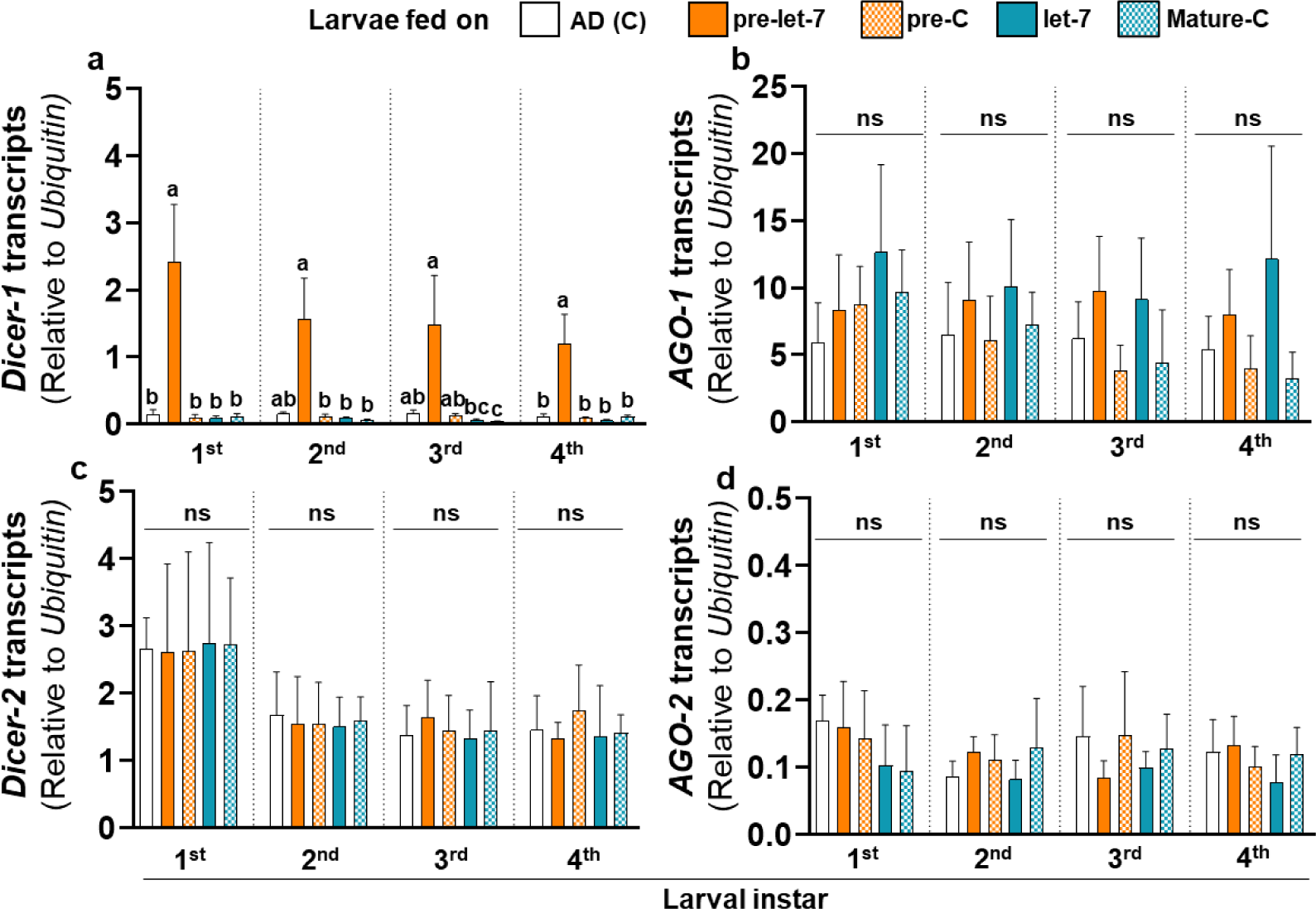
Effects of pre-let-7 and let-7 ingestion on the smRNA pathway components. Transcripts (relative to *Ubiquitin*) of miRNA pathway genes **a** *Dicer-1*, **b** AGO-1, and siRNA pathway genes **c** *Dicer-2* and **d** AGO-2. Significant differences in **a** were determined by the Kruskal-Wallis test and Dunn’s post hoc test (*p≤* 0.05), and in (**b**), (**c**), and (**d**) by one-way ANOVA and Tukey’s *post hoc* test (*p≤* 0.05). Letters a, b, and c denote differences between the larvae of the same instar feeding on different diets.

*Dicer-2* (Fig. 6b), *AGO-1* (Fig. 6c), and *AGO-2* (Fig. 6d) transcript levels did not vary upon pre-let-7 and let-7 ingestion; moreover, their levels did not correlate with the pre-let-7 or let-7 levels.

## Discussion

Interest in miRNA’s application in pest management is rapidly increasing due to miRNA’s target specificity, biodegradability, and conservation (Christiaens et al. 2020; Kumar et al. 2012; Liu et al. 2020). Several studies showed miRNA’s potential. However, developing a protocol based on those is difficult because the model species (Li et al. 2023; Zafar et al. 2021), miRNAs (Liu et al. 2022; Wen et al. 2021), miRNA conformations (pre- and ds mature) (Bally et al. 2020; Yogindran and Rajam 2021), miRNA lengths, and administration methods used in different studies were different. Furthermore, several reports did not provide information on instar-wise effects and, more importantly, phenotypic effects. It is natural because every study’s focus was different. Therefore, comparing pre- and mature miRNA’s effects across all the instars, vital for developing a ready-to-use method, is complicated based on the available information. We focused on comparing the efficacies and effects of the trophically delivered pre- and mature miRNAs. We systematically used one species, *P*. *xylostella*, one miRNA-let-7, and one administration method of miRNA feeding via AD. We started by optimizing the trophic delivery concentrations. We found that pre-let-7 and let-7’s 10 µg/g AD concentrations increased the endogenous pre-let-7 and let-7 levels and effectively downregulated the target genes. This was congruent with Whyard et al.’s, which showed 11 µg/g dsRNA concentration was effective against another lepidopteran *Manduca sexta* L. (Lepidoptera: Sphingidae) (2009).

Endogenous pre-let-7 and let-7 level elevation upon the miRNA ingestion indicated their uptake by the midgut cells form lumen; *Dicer-1* induction upon pre-let-7 ingestion suggested pre-let-7’s uptake and admission in the miRNA processing pathway. Since the *Dicer-1* induced and *AGO-1* and *AGO-2* did not, the smRNA is likely more tuned to perceiving the hairpin conformation pre-let-7 than let-7 in the trophic delivery system. Moreover, it raises the question of whether the exogenous mature ds miRNA has direct access to the cytoplasmic RISC or whether this access is *Dicer-1* dependent. Mature ds form is shorter than the pre-form; so, considering the high costs of RNA synthesis, it is more economical than the pre-. However, that (1) the mature ds miRNA’s higher molar concentration is required than the pre-miRNA for the target downregulation, (2) the *Dicer-1* induction by pre-miRNA indicates pre-miRNA’s interaction with the miRNA pathway, and (3) the pre-miRNA hairpins show more thermodynamic stability than the linear dsRNA (Bonnet et al. 2004) assert pre-miRNA as a more reliable option than the mature ds miRNA. Whether *Dicer-2* and *AGO-2*’s non-induction indicate the siRNA pathway’s inactiveness? If yes, then whether the cells perceive the trophically acquired mature ds miRNA as siRNA and whether this perception is the cause of its lower efficiency are good questions for future research on efficacy improvement and cost reduction.

Feeding pre-let-7 and let-7 downregulated the target transcripts *FTZ-F1* and *E74* in all four instars, confirming let-7’s post-transcriptional mode of action. Feeding pre-C and mature-C neither led to *Dicer-1* induction nor target suppression, indicating high miRNA-specificity of the miRNA pathway machinery and sequence-specificity in the target suppression. It also denoted that the hairpin cassette played no role in the target downregulation. Continuous pre-let-7 and let-7 ingestion caused larval mass and instar duration reduction and mortality increase in all instars compared to controls. Previous studies assessing the miRNA effects on larval mortality observed mortality only in specific instars (Gong et al. 2013; Gong et al. 2011; Ling et al. 2014; Sun et al. 2019). Similarly, in our instar-wise analysis, the first three instars showed significant mortality compared to controls. Although the mortality did not increase in the fourth instar, the miRNA-fed larvae showed lower mass and pupal eclosion than the controls. Finding such stark phenotypic effects in lepidopteran larvae is uncommon. This success could be attributed to our selection of let-7 as a candidate miRNA based on its prominent role in larval development.

Although let-7, with its stark effects, eased the miRNA administration standardization, let-7 cannot be recommended as a pest control miRNA. It is highly conserved across various insect taxa, implying that it will have low target taxon specificity and high off-target effects. The off-target effects on beneficial insects like pests’ natural enemies and pollinators are unwelcome in pest management. Therefore, our miRNA administration standardization using the let-7 candidate can help test the other natural and synthetic miRNAs’ potential in pest control.

Together, this work provides a proof-of-concept for the miRNA trophic delivery for pest control. The systematic investigation provided the optimum delivery concentrations and times. Pre-let-7 ingestion induced *Dicer-1* and effected the target gene silencing at a lower molar concentration than let-7; both pre-let-7 and let-7 caused the target gene downregulation after the same ingestion period (45-60 h). Although we used *P. xylostella* as a model and let-7 as a candidate miRNA, the miRNA administration standardization framework provided by this study will help discover different pest control-purpose miRNAs for the other pest systems.

## Author contribution

S.P. conceived the project. R.B. and S.P. designed and performed experiments, collected data. R.B. conducted statistical analyses. Both authors interpreted and discussed the results and wrote the manuscript. S.P. supervised the research.

## Supplementary Information

The online version contains supplementary material available at https://doi.org/-----.

## Acknowledgements

S.P. thanks the Department of Biotechnology, Government of India for the Ramalingaswami re-entry fellowship. R.B. thanks Indian Institute of Science Education and Research for the PhD fellowship. Authors thank, Mr. G. Pawar for help in insect culture maintenance and Dr. Gauri Binayak, Dr. Manish Kumar, and Ms. Kaushiki Kandalgaonkar for comments on the manuscript.

