## Supplementary information for "Trophic microRNA: Precursor and mature microRNA ingestion downregulates the target transcripts and hampers larval development in *Plutella xylostella*"

**Sagar Pandit**

Department of Biology, Indian Institute of Science Education and Research, Pune- 411008, Maharashtra, India

**Supplementary information**

**Fig. S1** Endogenous pre-let-7 and let-7 levels.

**Fig. S2** miRNA stability in AD.

**Table S1**. Primers used for the cDNA synthesis and quantitative real-time polymerase chain reactions (qPCRs).

**Table S2.** DNA templates used for *in vitro* transcription.

**Table S3.** Sequences of miRNAs supplemented in AD.

**Table S4.** Sequences of internal reference RNAs used in the AD-complemented pre-let-7 an dlet-7’s stability analysis.

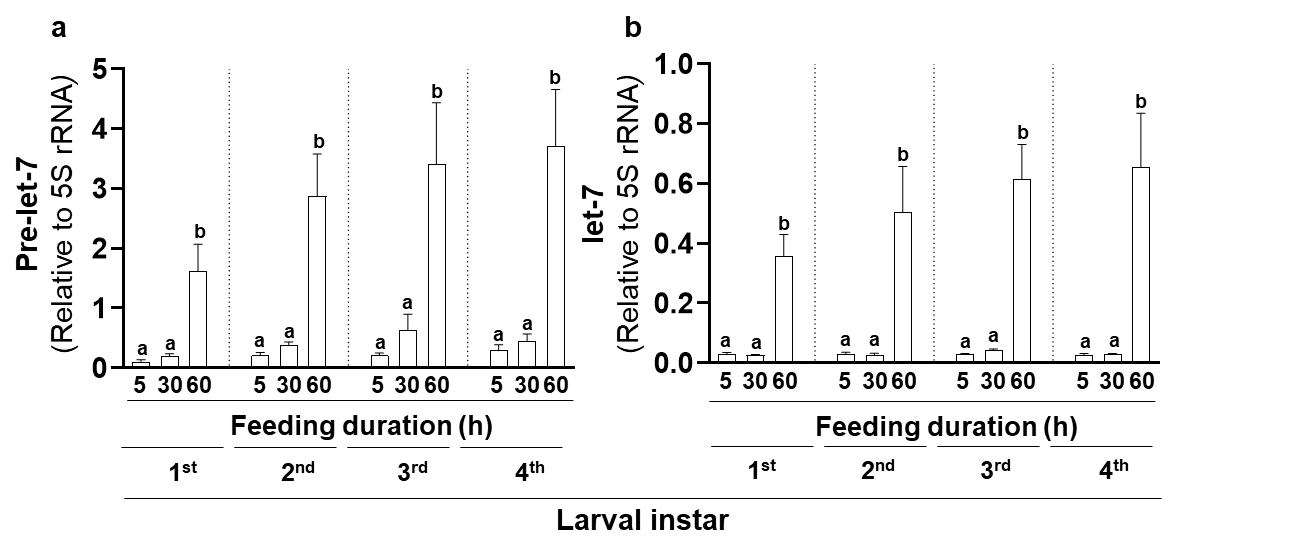

**Fig. S1** Endogenous pre-let-7 and let-7 levels. **a** Pre-let-7 (relative to *Ubiquitin*) and **b** let-7 (relative to 5s rRNA) levels in the early (5 h), mid (30 h), and late (pre-molting; 60 h) aged larvae of four larval instars. Significant differences were determined by the Kruskal-Wallis test and Dunn's *post hoc* test (*p≤* 0.05). Letters a, b, and c denote differences between the larvae of the same instar feeding on different diets.

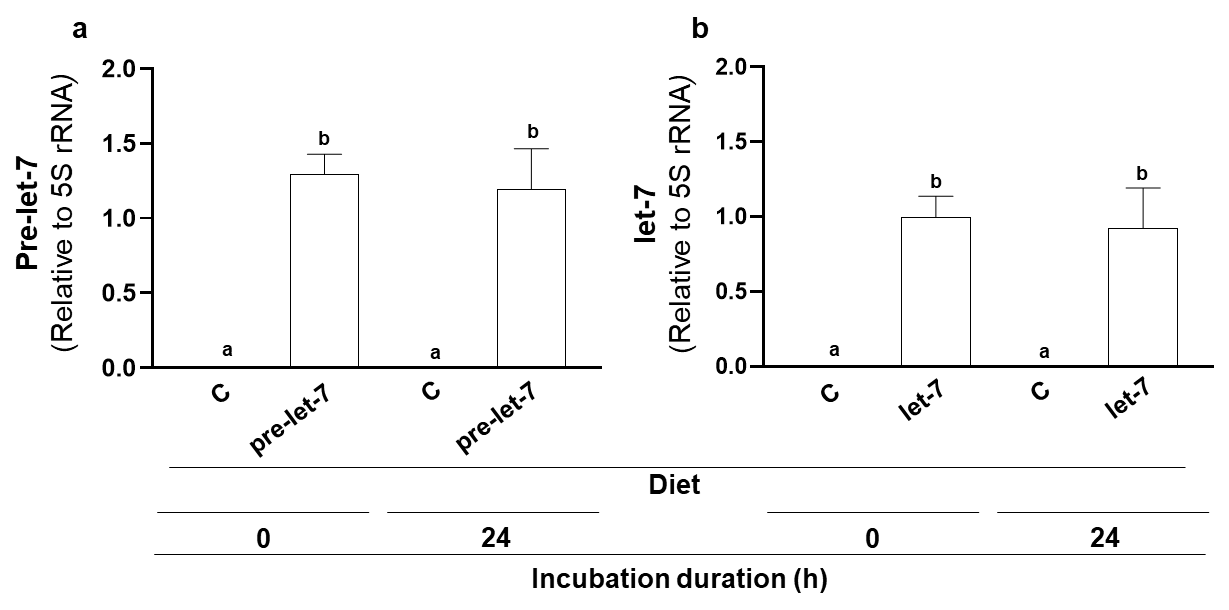

**Fig. S2** miRNA stability in AD. **a** Pre-let-7 (relative to *Ubiquitin*) and **b** let-7 (relative to 5s rRNA) transcripts in AD after 0 h and 24 h incubation. Significant differences were determined by the Kruskal-Wallis test and Dunn's *post hoc* test (*p≤* 0.05).

**Table S1**. Primers used for the cDNA synthesis and quantitative real-time polymerase chain reactions (qPCRs).

| **Transcript** | **Primer** | **Primer sequence** | **Used for** |
| --- | --- | --- | --- |
| Let-7 | Stem-loop RT | CTCAACTGGTGTCGTGGAGTCGGCAATTCAGTTGAGAACTATAC | cDNA synthesis |
| 5S rRNA | Stem-loop RT | GTCGTATCCAGTGCAGGGTCCGAGGTATTCGCACTGGATACGACAGCCAACG | cDNA synthesis |
| Let-7 | Forward | CCAGCTGGGTGAGGTAGTAGGTTGT | qPCR |
|  | Reverse | CTGGTGTCGTGGAGTCGGCAATT | qPCR |
| 5S rRNA | Forward | ACTGGATACGACAGCCAACG | qPCR |
|  | Reverse | GGCGTGGTCAGTACTTGGAT | qPCR |
| Pre-let-7 | Forward | GCCAGCTGAGGTAGTAGGTT | qPCR |
|  | Reverse | ACTGGCCAGGAAAGTTAGCA | qPCR |
| *FTZ-F1* | Forward | CAAACTGGCCTCATGTCCCT | qPCR |
|  | Reverse | TGGTAAGCTGCTCCTTGTGG | qPCR |
| *E74* | Forward | TCGACCTCTCCAATTGCCTG | qPCR |
|  | Reverse | CGACAGTTCCCCGAGAGATG | qPCR |
| *Dicer-1* | Forward | TTGCTGCATGAGGACGCTTA | qPCR |
|  | Reverse | TGAGGTCGTATGCCACACAC | qPCR |
| *Dicer-2* | Forward | TGATCTTCGACGAGTGCCAC | qPCR |
|  | Reverse | CTATGGTGGCGTGGAAGGTT | qPCR |
| *AGO1* | Forward | CTCCATGATGTACACCCCGG | qPCR |
|  | Reverse | GCTGGGCTTTGTAAAACGCA | qPCR |
| *AGO2* | Forward | GGAACTACTCTGAACCCGCC | qPCR |
|  | Reverse | GATGAAAGGCCGGGAAGTGA | qPCR |
| *Ubiquitin* | Forward | CGTGAAGACCCTTACTGGCA | qPCR |
|  | Reverse | TAGTCAGACAATGTGCGGCC | qPCR |

**Table S2.** DNA templates used for *in vitro* transcription.

| **Template** | **Sequence** |
| --- | --- |
| T7 promoter | TAATACGACTCACTATAGG |
| let-7 5p strand | CTATACAACCTACTACCTCACCTATAGTGAGTCGTATTA |
| let-7 3p strand | GGAAAGTTAGCAGGCTATACAGCCTATAGTGAGTCGTATTA |
| Scrambled let-7 5p strand | CTATACAACCTTCCCATCTACCTATAGTGAGTCGTATTA |
| Scrambled let-7 3p strand | GGAAGAAGAGCAGGCTATACAGCCTATAGTGAGTCGTATTA |
| pre-let-7 | ACTGGCCAGGAAAGTTAGCAGGCTATACAGTCCCCCGTTCGGTGTAATACTGTACTATACAACCTACTACCTCAGCTGGCCGAGCCTATAGTGAGTCGTATTA |
| scrambled pre-let-7 | ACTGGCCAGGAAGAAGAGCAGGCTATACAGTCCCCCGTTCGGTGTAATACTGTACTATACAACCTTCCCATCTAGCTGGCCGAGCCTATAGTGAGTCGTATTA |

**Table S3.** Sequences of miRNAs supplemented in AD.

| **miRNA** | **strand** | **Sequence** |
| --- | --- | --- |
| let-7 | 5p | UGAGGUAGUAGGUUGUAUAG |
|  | 3p | CUGUAUAGCCUGCUAACUUUCC |
| Mature-C (scrambled seed region) | 5p | UAGAUGGGAAGGUUGUAUAG |
|  | 3p | CUGUAUAGCCUGCUCUUCUUCC |
| Pre-let-7 | - | CUCGGCCAGCUGAGGUAGUAGGUUGUAUAGUACAGUAUUACACCGAACGGGGGACUGUAUAGCCUGCUAACUUUCCUGGCCAGU |
| Pre-C  (scrambled seed region in pre-let-7) | - | CUCGGCCAGCUAGAUGGGAAGGUUGUAUAGUACAGUAUUACACCGAACGGGGGACUGUAUAGCCUGCUCUUCUUCCUGGCCAGU |
| Yellow highlighted: Seed sequence  Green highlighted: Scrambled seed sequence | | |

**Table S4.** Sequences of internal reference RNAs used in the AD-complemented pre-let-7 an dlet-7’s stability analysis.

| **Internal reference RNA** | **Sequence** |
| --- | --- |
| 5S rRNA | ACTGGATACGACAGCCAACGACACGTGGTGTTCCCAGGCGGTCACCCATCCAAGTACTGACCACGCC |
| *Ubiquitin* | CGTGAAGACCCTTACTGGCAAGACAATCACTCTGGAAGTTGAAGCTTCAGACACCATCGAGAATGTGAAGGCCAAGATCCAAGACAAGGAAGGCATCCCTCCCGACCAGCAGCGACTTATCTTCGCTGGTAAACAACTAGAAGACGGCCGCACATTGTCTGACTA |
